# SERCA2b loss of function drives pigmentation by inducing adaptive ER stress and enhancing mitochondrial calcium uptake: significance in pathological hyperpigmentation associated with Darier’s Disease

**DOI:** 10.64898/2026.06.22.733702

**Authors:** Akshay Sharma, Suman Saurav, Preeti Sharma, Anushka Agrawal, Nutan Sharma, Gyan Ranjan, Divya Bhalla, Deepika Pandhi, Vamsi K Yenamandra, Jyoti Tanwar, Rajender K Motiani

## Abstract

Pigmentation is a critical protective mechanism that safeguards the skin against UV-induced damage, whereas dysregulated pigmentation predisposes to pigmentary disorders and skin malignancies. Although calcium signaling has emerged as an important regulator of melanogenesis, the identity of the calcium-handling proteins and the molecular mechanisms linking calcium dynamics to pigmentation remain poorly understood. Here, we identify the ER calcium pump SERCA2b as a negative regulator of pigmentation through modulation of ER stress and mitochondrial calcium uptake. We demonstrate that SERCA2b expression inversely correlates with pigmentation levels, and gain- and loss-of-function studies establish SERCA2b as a suppressor of melanogenesis. Mechanistically, SERCA2b depletion induces adaptive ER stress, enhances ER–mitochondrial proximity, and promotes mitochondrial calcium uptake. Notably, mutations in SERCA2b are associated with Darier disease, a condition characterized by hyperpigmented skin lesions, although the underlying mechanism remains unknown. To address this, we generated SERCA2b mutants corresponding to variants identified in Indian Darier’s disease patients and examined their effects on pigmentation, ER stress, and mitochondrial calcium dynamics. The mutant phenotypes closely recapitulated SERCA2b loss-of-function effects, demonstrating that adaptive ER stress and enhanced mitochondrial calcium signaling underlie hyperpigmentation associated with Darier’s disease. Importantly, treatment with 4-phenylbutyrate (4-PBA), an FDA-approved ER stress alleviator, rescued mutant-induced hyperpigmentation, reduced ER stress, and normalized mitochondrial calcium uptake. Collectively, our findings uncover a previously unrecognized role of SERCA2b in skin pigmentation, establish a mechanistic link between SERCA2b mutations and hyperpigmentation, and identify adaptive ER stress pathways as potential therapeutic target for pigmentary disorders.

## Introduction

Pigmentation is a defense phenomenon that protects skin from UV-rays induced DNA damage. Pigmentation is an outcome of melanin synthesis (melanogenesis) within a specialized lysosome related organelle, melanosome. Calcium (Ca^2+^) signaling is emerging as a critical regulator of pigmentation (Ahuja et al., 2024; Bellono & Oancea, 2014; Motiani et al., 2018; Saurav et al., 2025; Tanwar et al., 2023, 2024; Tanwar, Saurav, et al., 2022; Tanwar, Sharma, et al., 2022). Our lab has previously demonstrated a critical role of Endoplasmic Reticulum (ER) Ca^2+^ sensor Stromal Interaction Molecule 1 (STIM1) in driving physiological melanogenesis (Motiani et al., 2018; Tanwar, Sharma, et al., 2022). We showed that physiological melanogenic stimuli alpha-Melanocyte Stimulating Hormone (αMSH) induces ER Ca^2+^ release resulting in STIM1 activation. STIM1 in turn enhances cAMP production and enhanced transcription of melanogenic genes (Motiani et al., 2018). Further, our recent study has revealed role of mitochondrial calcium uniporter (MCU) in regulating melanogenesis by inducing expression of keratin filaments that in turn augments melanosome maturation and melanogenesis (Tanwar et al., 2023, 2024). While ER and mitochondrial Ca^2+^ signaling is emerging as a critical regulator of pigmentation, the explicit Ca^2+^ handling proteins that modulate pigmentation and downstream molecular mechanisms remain largely elusive.

Any perturbation in the skin pigmentation pathway predisposes to skin cancers and leads to pigmentary disorders. These disorders impart psychological trauma and severely effect the mental well-being of the patients. One such disorder is Darier’s disease that manifests patchy hyper-pigmentary phenotype along with other skin symptoms. Although Darier’s disease is associated with mutations in Sarcoplasmic/Endoplasmic Reticulum Calcium ATPase 2b (SERCA2b), the molecular mechanisms that drive hyper-pigmentary phenotype remain poorly understood.

We here reveal that SERCA2b acts as a negative regulator of pigmentation. Using several cellular models, we demonstrate that SERCA2b loss of function leads to increase in pigmentation while SERCA2b gain of function results in reduction in pigmentation. Further, we show that SERCA2b silencing induces adaptive ER stress and enhances mitochondrial Ca^2+^ uptake. Importantly, overexpression of Darier’s disease associated SERCA2b human mutation recapitulate SERCA2b silencing data suggesting that adaptive ER stress and elevated mitochondrial Ca^2+^ signaling could contribute to hyperpigmentation observed in Darier’s disease patients. Indeed, alleviation of ER stress with an FDA approved drug 4PBA rescues not only ER stress but also decreases mitochondrial Ca^2+^ uptake and pigmentation. Taken together, we reveal SERCA2b as a critical regulator of pigmentation by virtue of its role in regulating adaptive ER stress and mitochondrial Ca^2+^ signaling. Furthermore, we have revealed the potential of targeting adaptive ER stress pathway to modulate pigmentation.

## Materials and Methods

### Whole Exome Sequencing

A 10-year-old female presented with a provisional diagnosis of Darier disease. Post informed consent, genomic DNA was isolated from whole blood samples of the patient and family members. Whole Exome Sequencing (WES) was performed using illumina Nextera DNA library preparation kit, on Illumina HiSeq2500 sequencing platform following manufacturer’s protocol. The obtained quality passed paired-end sequences were aligned to the human reference genome (GRCh38) and variants were called using DRAGEN^TM^ software (Illumina Inc., USA). ANNOVAR was used for variant annotation based on variant’s position within the gene and the predicted impact on its protein function (Wang et al., 2010). Allelic frequencies in population datasets (1000Genome, gnomAD and IndiGen) allowed exclusion of common variants (Auton et al., 2015; *GnomAD*, n.d.; Jain et al., 2021). Clinically relevant variants (non-synonymous single nucleotide variants and small indels in the coding regions along with splice-site variants) were interpreted based on in-silico mutation prediction softwares (SIFT, Polyphen-2, MutationTaster2, CADD), existing variant databases (CLINVAR, dbSNP, HGMD) and literature evidence following American College of Genetics and Genomics and the Association for Molecular Pathology (ACMG-AMP) guidelines (Richards et al., 2015). Further, Sanger sequencing was used for validation and family segregation analysis of prioritized variants.

### Cell Culture

B16-F10 murine melanoma cells were procured from the American Type Culture Collection and cultured in Dulbecco’s Modified Eagle’s Medium-high glucose (DMEM-HG; Sigma-Aldrich, D5648) supplemented with 10% fetal bovine serum (FBS; Gibco, 10270106), 1X Antibiotic Antimycotic (Thermo Fisher Scientific, 15240062). The B16-F10 cells were maintained at 60-80% confluency in a humidified incubator with 5% CO_2_ levels.

Additionally, Human Epidermal Melanocytes, neonatal, lightly pigmented donor (HEMn-LP; Gibco, C0025C) were cultured in Medium 254 (M254; Gibco, M254CF) supplemented with human melanocyte growth supplement-2 (HMGS-2; Gibco, S0165) 1X Antibiotic Antimycotic (Thermo Fisher Scientific, 15240062) under the same environmental conditions. Experiments were conducted using cells between passages 3 to 6. Essential cell culture reagents used during experiments were phosphate-buffered saline, pH 7.2 (PBS; HIMEDIA, M1452), Trypsin (2.5%), no phenol red (Gibco, 15090046).

### LD pigmentation model

The Low-Density (LD) pigmentation-oscillator model was described earlier (Motiani et al., 2018; Tanwar et al., 2023, 2024; Tanwar, Sharma, et al., 2022). Briefly, B16 cells were seeded at 100 cells/cm^2^ in DMEM supplemented with 10% Fetal Bovine Serum in T75 flask and allowed to spontaneously pigment over 6-7 days.

### siRNA transfections

On the third day of the LD pigmentation model, siRNA transfections were conducted in T75 flasks. Each flask received 100nM of siRNA (sourced from Dharmacon) at a 1:3 (v/v) ratio with Dharmafect transfection reagent. siRNA and transfection reagent were mixed and incubated over the cells in OptiMEM (Gibco,31985070) media for 4–6hrs.

siRNA transfections were performed in B16 high density cells using siRNAs from Dharmacon. Dharmafect was used for transfecting B16 cells. siRNA and transfection reagent were mixed and incubated over the cells in OptiMEM media for 4–6hrs. For HEMn-LP, the cells were first trypsinized, and a cell pellet containing 7-10 x 10^5 cells was obtained. The cells were then resuspended in 100μL of the Nucleofector Solution for Human Melanocytes – Neonatal, provided in the Human Melanocyte-Neonatal Nucleofector Kit. (Lonza, VPD1003, U-024). Subsequently, 5μM siRNA (see Table 1) was added to this cell suspension. Media was changed 24hrs post transfection. Cells were harvested after 72hrs of transfection to capture phenotypic changes. The siRNAs were procured from Dharmacon.

**Table 1:** List of siRNAs.

| siRNA | Catalog Number |
| --- | --- |
| siNT | D-001810-10-20 |
| siSERCA2b (Mouse) | L-040968-00-0005 |
| siSERCA2b (Human) | L-004082-00-0005 |
| siMITF (Mouse) | L-047441-00-0010 |

The catalog numbers of siRNAs used in the study are provided in **Table 1** below:

### Generation of hSERCA2b overexpression construct

Human SERCA2b overexpression plasmid was generated by sub cloning human SERCA2b cDNA into mCherryC1 vector at SacI/KpnI sites. hSERCA2b cDNA was PCR amplified using Human SERCA2b (pcDNA 3.1+) (Addgene#75188) as template using Phusion High Fidelity Polyemrase (F503, Thermo). The amplified PCR product underwent restriction digestion with SacI/KpnI (NEB). Concurrently, the mCherryC1 empty vector was digested and treated with shrimp alkaline phosphatase (NEB).This was followed by gel purification of digested vector and PCR product. Vector and insert ligation was performed using Rapid DNA ligation Kit (K1422, Thermo). The ligation mix was transformed into chemically competent E. coli DH5α cells followed by plating onto Kanamycin containing agar plates. Positive clones were screened by restriction digestion analysis of plasmid DNA obtained by miniprep. Primers utilized for cloning are as follows:

hSERCA2bCDS FP

5’ – AAAGAGCTCAAATGGAGAACGCGCACACC-3’

hSERCA2bCDS RP

5’-GGGGGTACCAGACCAGAACATATCGCTAAAGTTAGTG-3’

### SERCA2b overexpression in B16 cells

B16 cells were seeded 48 h before transfection at a density of 5.0 × 10^4^ cells/well in 6-well plates. Human SERCA2b plasmid (1.5μg) was overexpressed in B16 cells plated at 60% confluency using Turbofect transfection reagent (R0532, Thermo) as per manufacturer’s protocol along with 1μm α-MSH treatment (Sigma-Aldrich, M4135) followed by cell termination after 48hrs of transfection. The overexpression was validated using RT-PCR and the effect of melanogenesis upon SERCA2b overexpression was analysed using Image J.

### Melanin content assay

Melanin-content assay was performed as described previously (Tanwar et al., 2024; Tanwar, Saurav, et al., 2022; Tanwar, Sharma, et al., 2022). The cells were lysed in 1N NaOH by heating at 80°C for 2 hrs and then absorbance was measured at 405nm. Melanin content was estimated by interpolating the sample readings on the standard melanin curve (μg/mL) obtained with synthetic melanin.

### Western blotting

Cells were lysed using NP40 lysis buffer supplemented with protease inhibitors. Typically,50-100μg proteins were subjected to SDS-PAGE (7.5–10%). Proteins from gels were then electro-transferred onto PVDF membranes. After blocking with 5% non-fat dry milk (NFDM) dissolved in Tris-buffered saline containing 0.1% Tween 20 (TTBS), blots were probed overnight at 4°C, with specific primary antibodies in TTBS containing 2% NFDM. The primary antibodies used were typically procured from Abcam, Cell signaling and were used at 1:500–1:2000 dilutions. The following day, membranes were incubated for 2hrs at room temperature with a horseradish-peroxidase-conjugated anti-mouse or anti-rabbit IgG antibody in TTBS containing 2% NFDM. Detection was performed using the enhanced chemiluminescence reagent (ECL Western blotting detection reagents; Amersham Biosciences). Quantification of bands was performed by densitometry using the ImageJ software. The catalogue number and company name for the antibodies are provided in the **Table 2** below:

**Table 2:** List of antibodies.

| Antibody | Company | Catalog Number |
| --- | --- | --- |
| DCT | Abcam | ab74073 |
| GP100 | Abcam | ab137078 |
| β-Tubulin | Abcam | ab21058 |
| SERCA2b | Abcam | ab183531 |

### Tyrosinase activity/DOPA assay

Tyrosinase-enzyme activity in cell lysates was assessed by performing DOPA assay as previously reported (Tanwar et al., 2024). Briefly, cell lysates were prepared in NP-40 lysis buffer and an equal amount of protein was run on a gel under nonreducing/native conditions. The gel was then immersed in phosphate buffer supplemented with tyrosinase substrate L DOPA (Sigma Chemicals, Bangalore, India). Enzyme activity corresponded to the formation of black color pigment.

### RT PCR

RT PCR was performed as described previously (Vashisht et al., 2018; Arora et al., 2021). mRNA extraction was performed with Qiagen RNeasy kit (Catalog#74106). cDNA was prepared using high-capacity cDNA reverse transcription kit from ThermoFisher (Waltham, MA, USA) (Catalog #4368814). Real-time PCR reactions were performed using SYBR green in Quant Studio 6 Flex from Applied Biosystems. The data were analyzed with Quant Studio real-time PCR software version 1.3. The expression of gene of interest was normalized to that of the housekeeping gene GAPDH. Primers were designed using Primer3 and checked by the NCBI Primer BLAST tool. Gene-specific primers were obtained from Eurofins. The nucleotide sequence of primers used in the study are provided in the below **Table 3**:

**Table 3:**
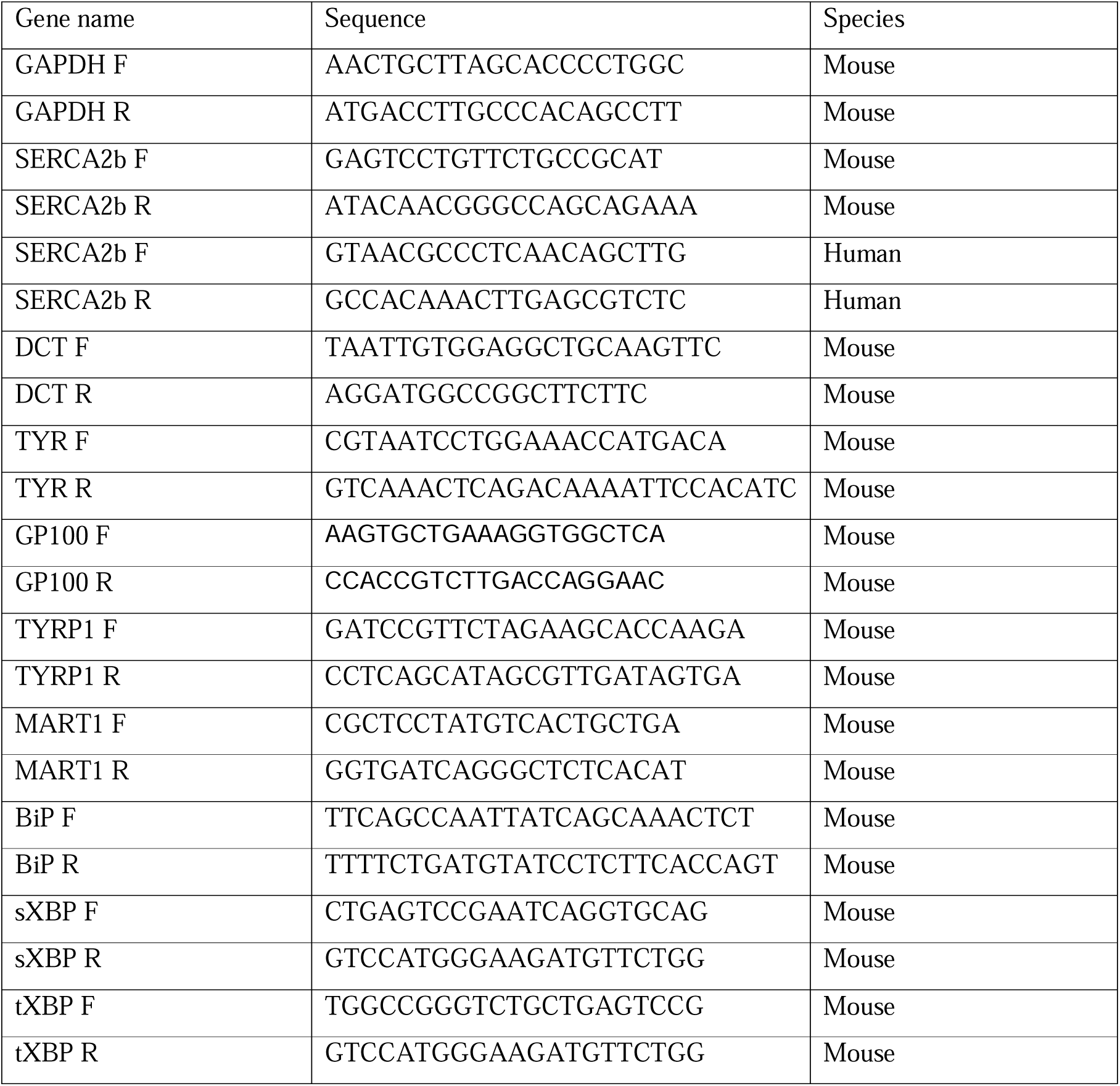
List of RT PCR primers.

**Table 4.**
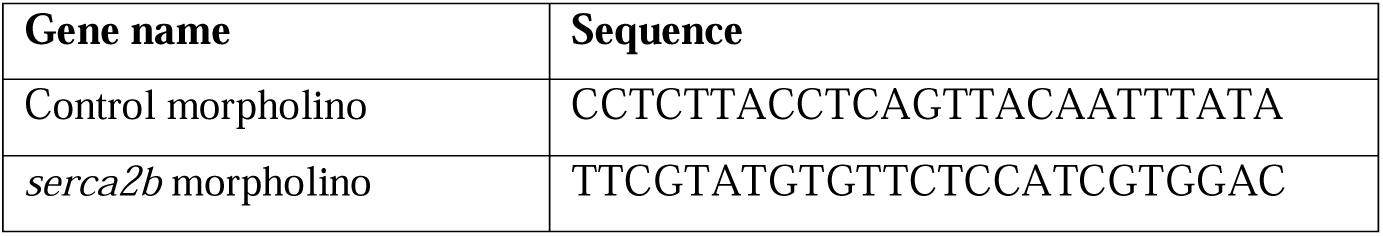
Details of morpholino used.

### Generation of mutant SERCA2b

SERCA2b C1484T (S495L) point mutant was generated through site directed mutagenesis using Q5 site directed mutagenesis kit (NEB, E0552S). Briefly, human SERCA2b overexpression plasmid (Addgene # 75188) was utilized as template for amplification by PCR using primers incorporating specific point mutation. The linear product was gel purified, then subjected to end phosphorylation, ligation, and subsequent Dpn1 digestion following the manufacturer’s protocol. The treated product was then transformed into chemically competent E. coli DH5α cells. Clones obtained post transformation were sequenced to identify presence of the specific mutation.

### SERCA2b Mutant overexpression in B16 cells

B16 cells were seeded 48hrs before transfection at a density of 5.0 × 10^4^ cells/well in 6-well plates. Human SERCA2b plasmid (1.5μg) and human SERCA2b mutant (1.5μg) was overexpressed in B16 cells plated at 60% confluency using Turbofect transfection reagent (R0532, Thermo) as per manufacturer’s protocol along with 1μM α-MSH treatment (Sigma-Aldrich, M4135) followed by cell termination after 48hrs of transfection. The overexpression was validated using RT-PCR and the effect of melanogenesis upon SERCA2b overexpression was analysed using Image J.

### Effect of SERCA2b Mutant OE upon α-MSH induced pigmentation in siSERCA2b condition

B16 cells were seeded 48hrs before transfection at a density of 15 × 10^3^ cells/well in 6-well plates. siRNA transfections in B16 cells plated using Lipofectamine 2000 (Invitrogen, 11668–019) transfection reagent as per manufacturer’s protocol. After 24 hrs of transfection, Human SERCA2b and Human SERCA2b Mutant (1.5μg) were overexpressed in transfected B16 cells using Turbofect transfection reagent (R0532, Thermo) as per manufacturer’s protocol along with 1μM α-MSH treatment (Sigma-Aldrich, M4135). The cells were terminated after 48hrs of over expression transfection. The pellet pictures were analysed using Image J.

### Effect of 4PBA upon α-MSH induced pigmentation in SERCA2b Mutant conditions

B16 cells were seeded 48hrs before transfection at a density of 5.0 × 10^4^ cells/well in 6-well plate. Human SERCA2b Mutant (1.5μg) was overexpressed in B16 cells plated at 60% confluency using Turbofect transfection reagent (R0532, Thermo) as per manufacturer’s protocol along with 1μM α-MSH treatment (Sigma-Aldrich, M4135) followed by addition of 4PBA after 6hrs. The cells were terminated after 48hrs of transfection. The pellet pictures were analysed using Image J.

### Cytosolic Ca^2+^ imaging

Live cell Ca^2+^ imaging was performed as reported previously (Raju et al., 2025). Cells were incubated at 37 °C for 30 min in a culture medium containing 4μM Fura-2 AM. After incubation, cells were washed 3 times and bathed in HEPES-buffered saline solution (140 mM NaCl, 1.13 mM MgCl2, 4.7 mM KCl, 2 mM CaCl2, 10 mM d-glucose, and 10 mM HEPES; pH 7.4) for ≥5 min before Ca^2+^ measurements were made. A digital fluorescence imaging system (Nikon Eclipse Ti2 microscope coupled with CoolLED pE-340 Fura light source and a high speed PCO camera) was used, and fluorescence images of multiple cells were captured and analyzed. Fura-2AM was excited alternately at 340 and 380 nm, and the emission signal was recorded at 510 nm. Figures showing Ca^2+^ traces are an average from several cells (the number of cells is denoted as “n” on each trace) on a single imaging dish. Each experiment was performed at least 3–4 times and the final data are plotted in the form of bar graphs.

### Mitochondrial Ca^2+^ imaging

Mitochondrial Ca^2+^ imaging was performed as reported previously (Tanwar et al., 2024). To perform mitochondrial Ca^2+^ imaging, B16 cells (0.5 x10 6 cells/well) were seeded onto confocal dishes. 24hours post plating (at 60% confluency), pCMV CEPIA2mt (Addgene plasmid # 58218) plasmid (1.5μg) was overexpressed using Lipofectamine 2000 (Invitrogen, 11668-019). After 24hrs post transfection, cells were washed 3 times and bathed in HEPES-buffered saline solution (140mM NaCl, 1.13mM MgCl2, 4.7mM KCl, 2mM CaCl2, 10mM D-glucose, and 10mM HEPES; pH 7.4) for 5min prior to Ca^2+^ measurements. The Nikon Eclipse Ti2 microscope was employed to capture and analyze fluorescence images of various cells using a 60X oil objective. The CEPIA2mt-transfected cells were excited at 488 nm, and the emission signal was captured at 500-550nm. A 100μM histamine in a Ca^2+-^free bath solution was used as a stimulus to release Ca^2+^ from the ER via IP3Rs.Experiments were performed at least 3 times and the final data are plotted in the form of violin plots where the number of cells is denoted as “n”. pCMV CEPIA2mt was a gift from Masamitsu Lino (Addgene plasmid # 58218).

### Zebrafish husbandry

Assam wild type (ASWT) (Lalwani et al., 2012) was used and were maintained and handled in compliance with the protocols and guidelines approved by the institutional animal ethic committee RCB. Special care was taken to ensure minimal distress to the animals.

### Microinjections

Antisense ATG targeting morpholino oligonucleotide (MO) was designed to knockdown the serca2b in zebrafish (GeneTools, USA). Using nucleus-free water, we resuspended the MO to a final stock concentration of 1mM. 3nl of 100uM of MO was injected into hindbrain ventricle of 2dpf zebrafish which were pretreated with (0.003%) PTU (N-phenylthiourea, Sigma Aldrich) and after injection was transferred to embryo water (E3). After 24hrs post injection (hpi) imaging was performed on Nikon SMZ800N. Quantification was done by measuring mean intensity subtracted by background and plotted using GraphPad v10. Human SERCA2b wild-type and SERCA 2b mutant c.1484 C<T was amplified and *in-vitro* transcribed (IVT) using T7 mMassage mMachine kit (Thermo scientific). 3nl of 50ng/uL of IVT product was injected into the single-cell stage ASWT embryos and transferred to embryo water (E3) and were incubated at 28.5□°C. Embryo water was changed every 24hrs and the dead and undeveloped embryos were removed simultaneously. Imaging was performed at 48hpf on Nikon SMZ800N and quantification was done by measuring mean intensity subtracted by background and plotted using GraphPad v10.

### Zebrafish 4PBA treatment

ASWT zebrafish embryos were treated 48 hours post fertilization (hpf) with 300μM 4PBA for 24 hrs.

### Measuring the distance between ER and Mitochondria

A split-GFP-based contact site sensor (SPLICS) was used to quantify the contact sites between the endoplasmic reticulum and mitochondria across a range of distances (Cieri et al., 2017). B16 cells were transfected with 1µg of the SPLICS Mt-ER Short P2A plasmid (Addgene, 164108) using the TurboFect transfection reagent at a 1:2 ratio (w/v). Live-cell imaging was performed 24hrs after the plasmid transfection, utilizing a confocal microscope equipped with a 63X/1.40 oil immersion objective **(Laser Scanning Confocal Microscope: LSM 880, Carl Zeiss)**. The cells were excited with 488 nm lasers, and the resulting fluorescence emissions were captured within the 497-572nm range. The confocal images were then deconvoluted using ZEN 2.3 SP1 FP1 (black) software (version 14.0) and analyzed using Fiji software.

### Effect of 4PBA upon **α**-MSH induced pigmentation

B16 cells were seeded at a density of 5.0 × 10^4^ cells/well in 6-well plates. 1μM α-MSH treatment (Sigma-Aldrich, M4135) was given, followed by the addition of 4PBA after 6hrs. The cells were terminated after 48hrs. The pellet pictures were analysed using Image J.

### Statistical Analysis

All statistical analysis was performed using GraphPad Prism 10 software. All experiments were performed at least 3 times. Data are presented as mean ± SEM and either one-sample or unpaired student’s t-test was performed for determining statistical significance between 2 experimental samples, whereas one-way ANOVA was performed for the comparison of 3 samples. A p-value < 0.05 was considered as significant and is presented as “*”; p-value < 0.01 is presented as “**”; p-value < 0.001 is presented as “***”and p-value < 0.0001 is presented as “****”.

## Results

### SERCA2b negatively regulates pigmentation *in vitro*

To explore the role of SERCA2b in pigmentation, we examined levels of SERCA2b in low-density culturing (LD) induced pigmentation. In the LD pigmentation model, B16 mouse melanoma cells are cultured at a low density of 100 cells/cm^2^ allowing them to pigment autonomously over a period of six-seven days (**Fig. 1A**) (Motiani et al., 2018; Tanwar et al., 2024; Tanwar, Sharma, et al., 2022).We observed that both mRNA and protein expression of SERCA2b decreases significantly with increase in pigmentation in B16 cells in the LD model (**Fig.1B-D)**.

**Figure 1.**
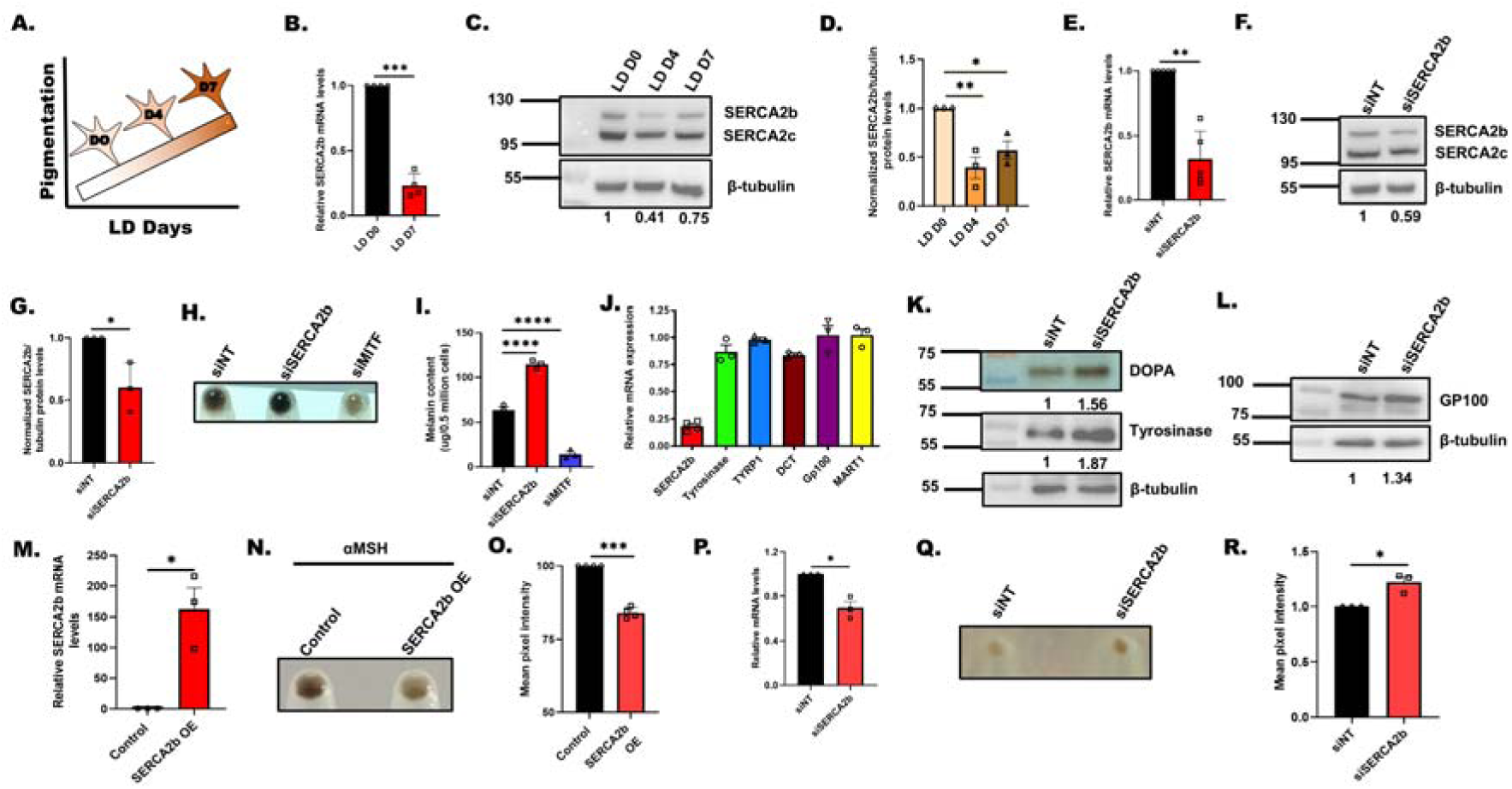
SERCA2b negatively regulates pigmentation. **(A)** Diagrammatic representation of B16 low density (LD) pigmentation model. **(B)** qRT–PCR analysis showing decrease in SERCA2b mRNA expression with pigmentation in B16 LD model (N=3). **(C)** Representative western blot showing decrease in SERCA2b protein expression on LD day 4 and 7 in comparison with LD day 0 (N=3). Densitometric analysis using ImageJ is presented below the blot. **(D)** Densitometric quantitation showing decrease in SERCA2b protein levels on LD day 4 and 7 in comparison with LD day 0 (N=3). **(E)** qRT–PCR analysis showing decrease in SERCA2b mRNA expression upon SERCA2b silencing in B16 LD D7 cells (N=5). **(F)** Representative western blot confirming siRNA based silencing of SERCA2b on LD day 7 B16 cells. Densitometric analysis using ImageJ is presented below the blot (N=3). **(G)** Densitometric quantitation showing SERCA2b levels on LD day 7 in siNT control and siSERCA2b condition (N=3). **(H)** Representative pellet images of siNT control, siSERCA2b and siMITF on LD day 7 (N=3). **(I)** Melanin content estimation of siNT, siSERCA2b and siMITF B16 cells on LD day 7 (N=3). **(J)** qRT–PCR analysis showing tyrosinase, TRP1, DCT, GP100 and MART1 mRNA expression upon SERCA2b silencing in B16 cells (N=3-4). **(K)** DOPA assay showing activity of tyrosinase enzyme and representative western blot for tyrosinase expression on LD day 7 upon SERCA2b silencing as compared to siNT control. Densitometric analysis using ImageJ is presented below the blot. **(L)** Representative western blot showing expression of GP100 on LD day 7 upon SERCA2b silencing as compared to siNT control. Densitometric analysis using ImageJ is presented below the blot. **(M)** qRT–PCR analysis showing increase in SERCA2b mRNA expression upon ectopic expression of SERCA2b in B16 high density (HD) cells (N=3). **(N)** Representative pellet images of mCherry empty vector (EV) control and SERCA2b overexpressing (OE) B16 high density (HD) cells upon αMSH treatment (N=4). **(O)** Quantitation of mean pixel intensity of control plasmid and SERCA2b overexpression upon αMSH treatment (N=4). **(P)** qRT–PCR analysis showing decrease in SERCA2b mRNA expression upon SERCA2b silencing in primary human melanocytes (N=3). **(Q)** Representative pellet images of siNT control, siSERCA2b in primary human melanocytes (N=3). **(R)** Quantitation of mean pixel intensity of siNT control, siSERCA2b in primary human melanocytes (N=3). Data presented are mean ± S.E.M. For statistical analysis, unpaired student’s t-test was performed for panel G, one sample t test for panels B, E, M, O, P, R and one-way ANOVA followed by Tukey’s post hoc test for panels D, I using Graphpad Prism software. (Here, * p <0.05; ** p < 0.01; ∗∗∗ p < 0.001 and ∗∗∗∗p < 0.0001)

We next investigated role of SERCA2b in pigmentation by performing loss of function and gain of function studies. Initially, we characterized the siRNA targeting SERCA2b in B16 cells, achieving approximately a 70% reduction in SERCA2b mRNA levels with the targeted siRNA. (**Fig. 1E**). We further validated the efficacy of the SERCA2b siRNA at protein levels and detected almost 40% decrease in SERCA2b levels (**Fig. 1F and G**). Next, we silenced SERCA2b in LD pigmentation model and observed a drastic increase in the pigmentation as evident from pellet picture (**Fig. 1H**). In this assay, we used MITF (microphthalmia-associated transcription factor) as a positive control and as expected MITF knockdown decreased pigmentation (**Fig. 1H**). We quantitated the changes in pigmentation by estimating melanin content under various conditions and found that SERCA2b silencing significantly enhances pigmentation (**Fig. 1I**). Further, we analyzed mRNA expression of key melanogenic enzymes and melanosome structural proteins i.e., tyrosinase, tyrosinase-related protein 1 (TYRP1), tyrosinase-related protein 2/Dopachrome Tautomerase (DCT), GP100 and MART1 in siNT and siSERCA2b cells. We observed that SERCA2b silencing did not alter the mRNA levels of key melanogenic enzymes and proteins (**Fig. 1J**).

We next examined the levels of key melanosome structural protein Gp100 and activity & expression of rate limiting enzyme in the melanogenesis process i.e. tyrosinase upon the silencing of SERCA2b. As presented (**Fig. 1K and L**), we observed a robust increase in Gp100 levels, tyrosinase activity and tyrosinase expression upon SECA2b knockdown. This data suggests that SERCA2b negatively regulates melanosome biogenesis and melanin synthesis.

Next, we performed SERCA2b gain of function experiments. We cloned SERCA2b in a mcherry construct and overexpressed it in B16 cells. We observed a drastic increase in SERCA2b levels upon ectopic expression of SERCA2b (**Fig. 1M**). We subsequently investigated the impact of SERCA2b overexpression on physiological pigmentation stimulated by alpha-melanocyte stimulating hormone (αMSH). Our findings revealed that this overexpression reduces αMSH-induced pigmentation. (**Fig. 1N**). We further quantitated these phenotypic changes and observed that the ectopic expression of SERCA2b leads to a significant reduction in αMSH-induced melanogenesis in comparison to empty vector control (**Fig. 1O**). Taken together, our results demonstrate that SERCA2b negatively regulates melanogenesis in B16 cells.

To further corroborate role of SERCA2b in pigmentation, we studied it in primary human melanocytes. To examine the role of SERCA2b in regulating melanogenesis, we silenced it using human siRNAs in primary human melanocytes. We validated SERCA2b knockdown by performing qRT-PCR. We observed around 40% decrease in SERCA2b mRNA levels in primary melanocytes transfected with siSERCA2b as compared to siNT (**Fig. 1P**). The knockdown of SERCA2b resulted in visible increase in melanogenesis as evident in the pellet images of primary melanocytes (**Fig. 1Q**). We quantitated the change in melanogenesis upon SERCA2b silencing in primary melanocytes using Image J based analysis and observed a significant increase in pigmentation in the siSERCA2b condition compared to the siNT control (**Fig. 1R**). Collectively, our data from B16 cells and primary human melanocytes show that SERCA2b is a critical negative regulator of pigmentation.

### SERCA2 regulates pigmentation *in vivo*

We next investigated the role of SERCA2b in pigmentation *in vivo* using zebrafish model system. We injected SERCA2b morpholinos into hindbrain ventricle of zebrafish at 2 dpf (days post fertilization) and then observed changes in pigmentation post injections (**Fig. 2A**). Phenotypically, we observed significant increase in the pigmented black-colored melanophores in SERCA2b morphants as compared to the control embryos at 24hpi (hours post injection) (**Fig. 2B**). We quantitated this increase in pigmentation by Image J based analysis (**Fig. 2C**). To further strengthen our *in vivo* data we performed gain of function studies in zebrafish. We injected human SERCA2b RNA in zebrafish and observed decrease in the pigmented melanophores as compared to the control embryos at 48 hpf (hours post fertilization) (**Fig. 2D**). We quantitated this decrease in pigmentation by Image J based analysis (**Fig. 2E**). Collectively, our zebrafish data demonstrate an important role of SERCA2b in regulating pigmentation in vivo.

**Figure 2.**
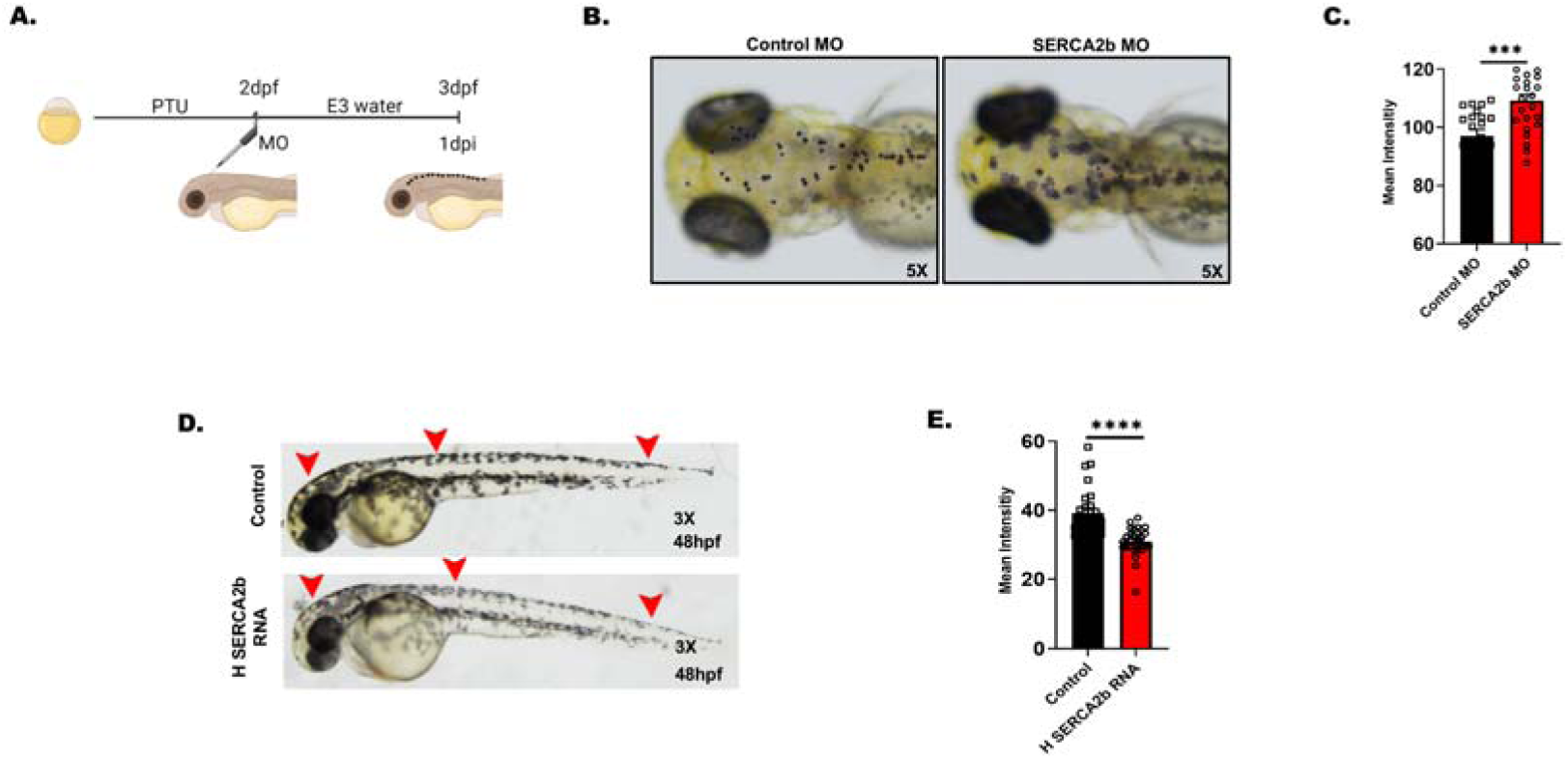
SERCA2b regulates pigmentation *in vivo* in zebrafish. **(A)** Schematic diagram showing MO injections into hindbrain ventricle of 2 dpf zebrafish. **(B)** Representative bright-field cranial images of wild-type zebrafish embryos injected with either control morpholino or SERCA2*b* morpholino at 24hpi (hours post injection). **(C)** Quantitation of mean pixel intensity of control MO and SERCA2*b* MO in zebrafish embryos post injection (each dot represents individual embryo from 3 independent experiments, N= ∼25). **(D)** Representative bright-field images of zebrafish embryos injected with either control or human *SERCA2b* RNA at 48hpf. **(E)** Quantitation of mean pixel intensity of zebrafish embryos injected with either control or human *SERCA2b* RNA (each dot represents individual embryo from 3 independent experiments, N= ∼30). Data presented are mean ± S.E.M. For statistical analysis, Mann Whitney test was performed for panels C, E using Graphpad Prism software. (Here, * p <0.05; ** p < 0.01; ∗∗∗ p < 0.001 and ∗∗∗∗p < 0.0001)

### SERCA2b silencing reduces ER Ca^2+^ levels and induces adaptive ER stress

To elucidate the molecular mechanism working downstream of SERCA2b to regulate pigmentation, we investigated the Ca^2+^ levels in the endoplasmic reticulum (ER), given that the primary cellular function of SERCA2 is to facilitate Ca^2+^ influx from cytosol into the endoplasmic reticulum (ER) (Stutzmann & Mattson, 2011). Therefore, we examined the ER Ca^2+^ stores upon SERCA2b knockdown. We utilized ionomycin to release ER Ca^2+^ stores into cytosol and used FURA2-AM (ratio-metric Ca^2+^ probe) to measure the ionomycin-induced increase in cytosolic Ca^2+^ levels, providing an estimate of Ca^2+^ stored in the ER. We noticed that siSERCA2b condition has significantly lower ER Ca^2+^ stores in comparison to siNT control (**Fig. 3. A and B**). Literature suggests that decrease in ER Ca^2+^ levels result in adaptive ER stress or unfolded protein response (UPR) (Chen et al., 2023; Makio et al., 2024). Hence, we investigated if adaptive ER stress is induced upon SERCA2b silencing. We examined mRNA levels of ER stress markers such as BiP, sXBP and tXBP and observed that SERCA2b silencing results in adaptive ER stress (**Fig. 3. C-E**).

**Figure 3.**
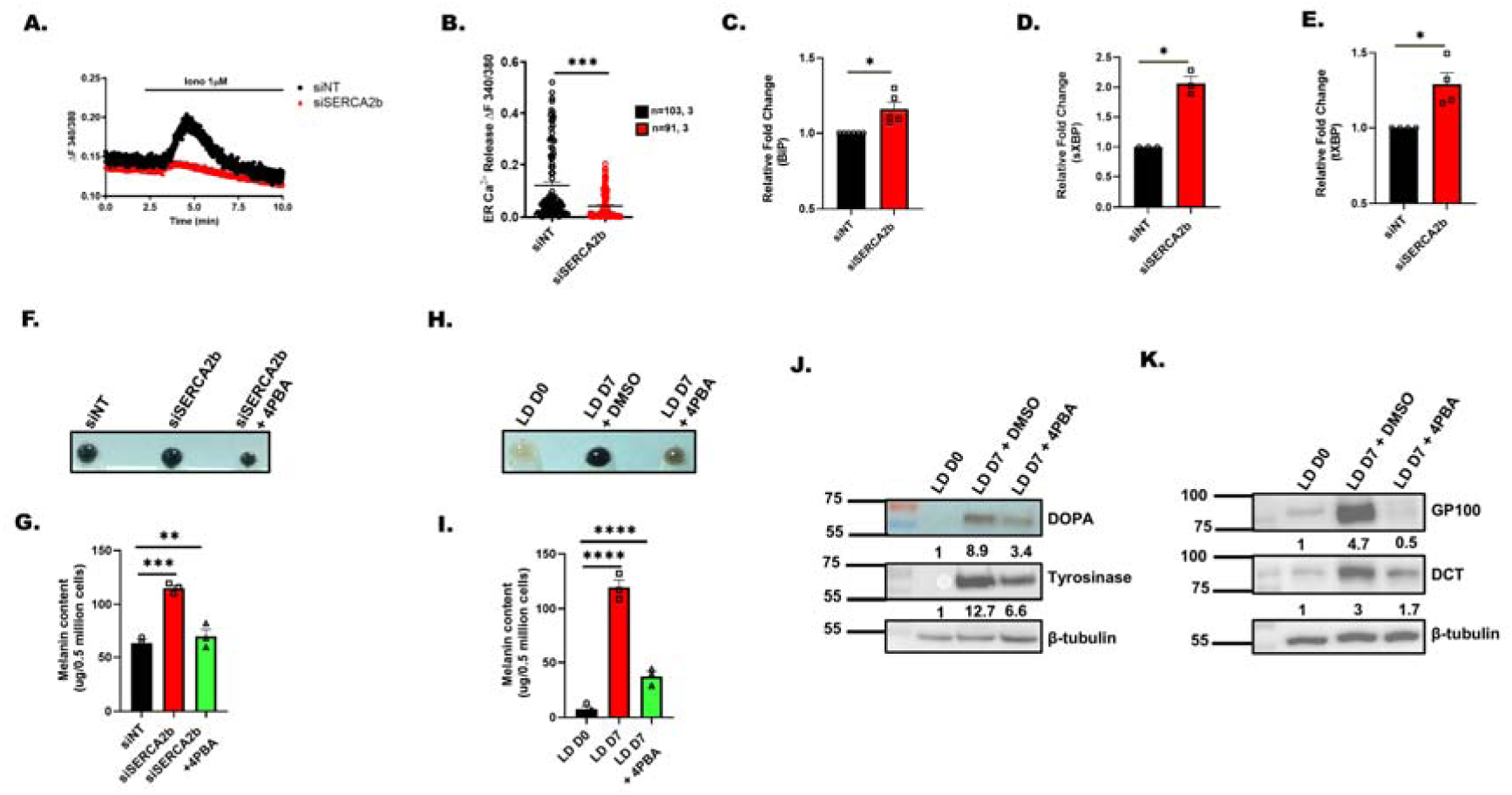
SERCA2b silencing reduces Ca^2+^ levels and induces adaptive ER stress. **(A)** Representative Ca^2+^ imaging traces of siNT and siSERCA2b where “n” denotes the number of cells in that particular trace stimulated with 1µM ionomycin in Ca^2+^-free buffer **(B)** The extent of SOCE was calculated from 103 siNT and 91 siSERCA2b B16 cells, which were imaged from several experiments/traces (“n□=□x, y” where “x” denotes total number of cells imaged and “y” denotes number of traces recorded). **(C)** qRT–PCR analysis showing increase in BiP mRNA expression upon SERCA2b silencing in B16 HD cells along with αMSH treatment (N=5). **(D)** qRT–PCR analysis showing increase in sXBP1 mRNA expression upon SERCA2b silencing in B16 HD cells along with αMSH treatment (N=3). **(E)** qRT–PCR analysis showing increase in tXBP1 mRNA expression upon SERCA2b silencing in B16 HD cells along with αMSH treatment (N=4). **(F)** Representative pellet images of siNT control, siSERCA2b and siSERCA2b+4PBA on LD day 7 (N=3). **(G)** Melanin content estimation of siNT control, siSERCA2b and siSERCA2b+4PBA on LD day 7 (N=3). **(H)** Representative pellet images of LD D0, LD D7+DMSO and LD D7+4PBA B16 cells (N=3). **(I)** Melanin content estimation of LD D0, LD D7+DMSO and LD D7+4PBA (N=3) B16 cells. **(J)** DOPA assay showing activity of tyrosinase enzyme and representative western blot for tyrosinase expression in LD D0, LD D7+DMSO and LD D7+4PBA B16 cells. Densitometric analysis using ImageJ is presented below the blot. **(K)** Representative western blot showing expression of GP100 and DCT in LD D0, LD D7+DMSO and LD D7+4PBA B16 cells. Densitometric analysis using ImageJ is presented below the blot. Data presented are mean ± S.E.M. For statistical analysis, unpaired student’s t-test was performed for panel B, one sample t test for panels C, D, E, and one-way ANOVA followed by Tukey’s post hoc test for panels G, I using Graphpad Prism software. (Here, * p <0.05; ** p < 0.01; ∗∗∗ p < 0.001 and ∗∗∗∗p < 0.0001)

### 4PBA rescues SERCA2 silencing induced pigmentation

Since SERCA2b knockdown resulted in induction of ER stress and enhanced pigmentation, we investigated whether ER stress plays a role in the pigmentation induced by SERCA2b silencing. We used a well-established FDA-approved ER stress alleviator 4-phenylbutyric acid (4-PBA) to rescue ER stress (Iannitti & Palmieri, 2011). Interestingly, 4-PBA almost completely rescues the increase in pigmentation observed upon SERCA2b knockdown thereby suggesting that downstream of SERCA2b silencing pigmentation is at least partially regulated by induction of ER stress (**Fig. 3. F and G**).

We next tested effect of 4-PBA in B16 LD pigmentation model. We observed decrease in pigmentation as evident in B16 LD day 7 cell pellets upon 4-PBA treatment as compared to vehicle control (**Fig. 3H**). We further quantitated these changes in pigmentation by estimating melanin content and found that 4-PBA treatment significantly decrease pigmentation (**Fig. 3I**).

We then examined the levels of key melanosome structural protein Gp100, expression of melanogenic enzyme (DCT and tyrosinase) and tyrosinase activity. We observed decrease in Gp100, DCT levels and tyrosinase activity as well as tyrosinase expression upon 4-PBA treatment. Collectively this data suggests that 4-PBA treatment rescues SERCA2b silencing induced pigmentation and decreases B16 LD pigmentation (**Fig. 3, J and K**).

We next surveyed the literature to identify how ER stress can drive pigmentation. Recently, it was reported that ER stress leads to increase mitochondria-endoplasmic reticulum contacts (MERCs) and enhanced mitochondrial Ca^2+^ uptake (Bassot et al., 2023). Interestingly, the recent work from our lab has demonstrated that mitochondrial Ca^2+^ uptake is a critical determinant of pigmentation (Tanwar et al., 2023, 2024). Therefore, we examined if downstream of SERCA2b silencing MERCs and mitochondrial Ca^2+^ uptake was affected.

### SERCA2b knockdown decreases ER-mitochondrial distance and augments mitochondrial Ca^2+^ uptake

We used split GFP-based contact site sensor SPLICS_(short)_ (ER-Mito) (Cieri et al., 2017) to measure ER-mitochondrial distance upon SERCA2b silencing (**Fig. 4 A**). We observed an increase in MERCs upon SERCA2b silencing compared to the siNT control **(Fig. 4, B and C)**. This data suggests that the ER and mitochondria come closer together, upon SERCA2b silencing. Next, we asked whether these increased ER-mitochondrial contacts could increase mitochondrial Ca^2+^ downstream of SERCA2b silencing.

**Figure 4.**
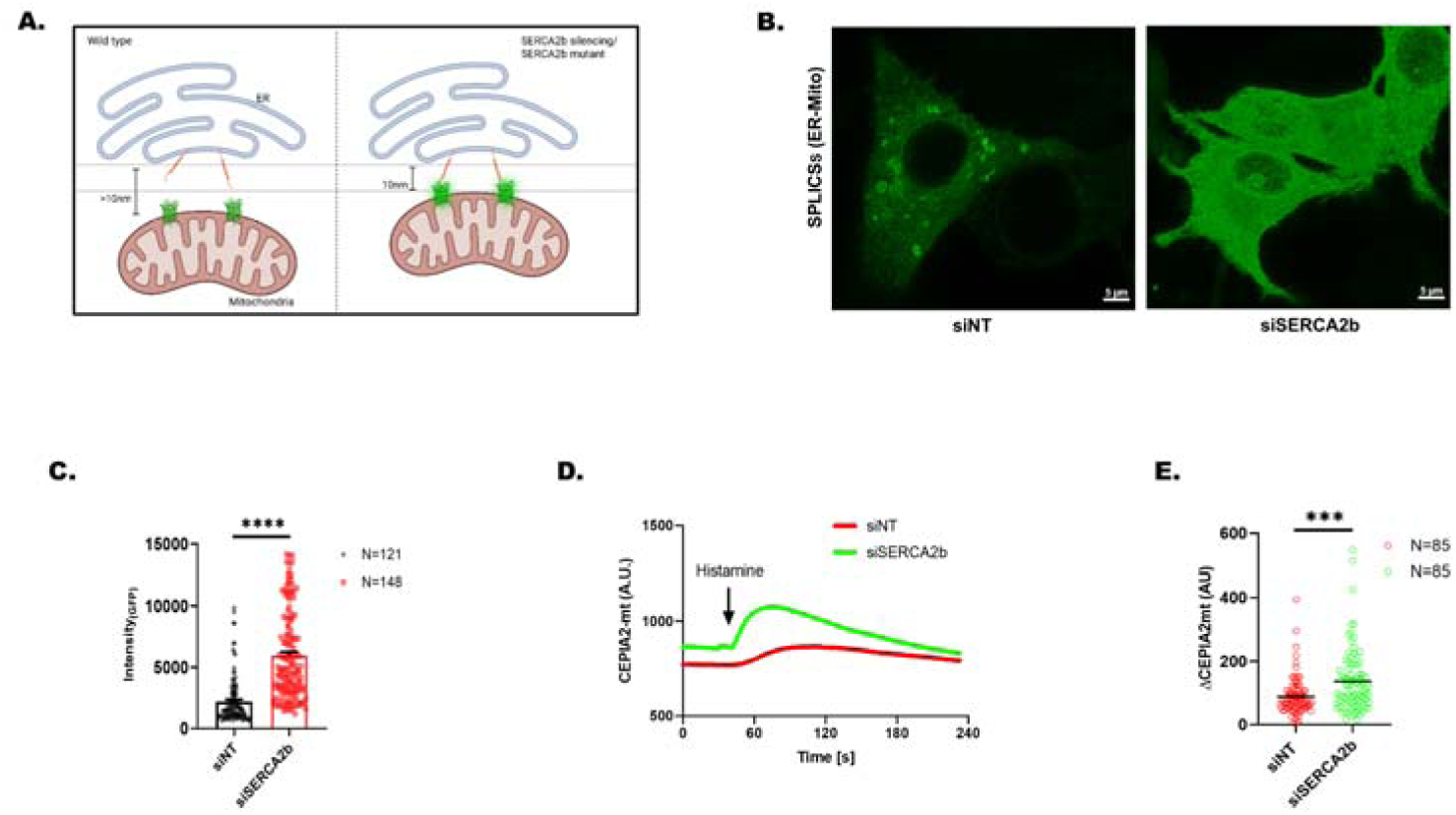
SERCA2b silencing regulates ER-mitochondrial distance and mitochondrial Ca^2+^ uptake. **(A)** A schematic illustration demonstrating SPLICS Mt-ER (ER-Mito) probes to measure distance between ER and mitochondria. **(B)** Representative images of confocal imaging in B16 cells transfected with siNT or siSERCA2b along with SPLICS Mt-ER Short P2A construct, scale bar, 5µm. **(C)** Bar graph showing the quantification of the GFP intensity, in B16 cells transfected with siNT or siSERCA2b along with SPLICS Mt-ER Short P2A construct, where ‘N’ denotes the number of cells imaged. **(D)** Representative mitochondrial Ca^2+^ imaging traces using the CEPIA2mt probe in siNon-Targeting (siNT) control and siSERCA2b B16 cells, stimulated with 100μM histamine. **(E)** Bar graph showing the quantification of Ca^2+^ imaging traces stimulated with 100μM histamine, where ‘N’ denotes the total number of regions of interest (ROI) in that trace. Data presented are mean ± S.E.M. For statistical analysis, unpaired student’s t-test was performed for panel C, E using Graphpad Prism software. (Here, * p <0.05; ** p < 0.01; ∗∗∗ p < 0.001 and ∗∗∗∗p < 0.0001)

We used genetically encoded Ca^2+-^measuring organelle-Entrapped Protein Indicators (CEPIA) for studying mitochondrial Ca^2+^ dynamics (Tanwar et al., 2024). We transfected B16 cells with a mitochondrial matrix specific CEPIA probe (CEPIA2mt) to analyzed mitochondrial Ca^2+^ uptake upon SERCA2b silencing. We used histamine, a physiological agonist, for studying mitochondrial Ca^2+^ uptake. Histamine releases Ca^2+^ from endoplasmic reticulum thereby enhancing cytosolic Ca^2+^ levels. This rise in cytosolic Ca^2+^ stimulates mitochondrial Ca^2+^ uptake via Mitochondrial Ca^2+^ Uniporter (MCU) complex (Tanwar et al., 2024). Our live cell mitochondrial Ca^2+^ imaging assays show that upon SERCA2b knockdown mitochondrial Ca^2+^ uptake is augmented (**Fig. 4D and E**). This increased mitochondrial Ca^2+^ uptake increases pigmentation upon SERCA2b silencing.

### Darier disease associated SERCA2b mutations recapitulate phenotype and lead to reduction in ER Ca^2+^

A 10-year-old patient, suspected to have Darier disease, was clinically characterized by multiple discrete, warts, hyperkeratotic papules predominantly involving the flexural regions, with overlying pustules, crusted erosions, and areas of hyperpigmentation **(Fig. 5A)**. The patient reported exacerbation of lesions during summer months. Nail examination revealed longitudinal erythronychia, while oral examination was not remarkable. Family history indicate that her mother and two younger siblings had similar lesions, whereas her father and two other siblings were apparently healthy. WES analysis revealed a previously reported heterozygous ‘pathogenic’ variant in the exon 12 of *ATP2A2* gene (NM_170665.4:c.1484C>T:p.Ser495Leu). This variant is absent in publicly available population databases like 1000 Genomes, gnomAD (Genome Aggregation database) and IndiGen and is predicted to be deleterious by computational prediction scores (SIFT: 0; PolyPhen: 0.9 and CADD Phred: 35). Sanger sequencing confirmed the presence of prioritized variant in heterozygous state in the proband, mother and affected siblings while the homozygous wild-type allele in father and remaining two siblings (**Fig. 5B**).

**Figure 5.**
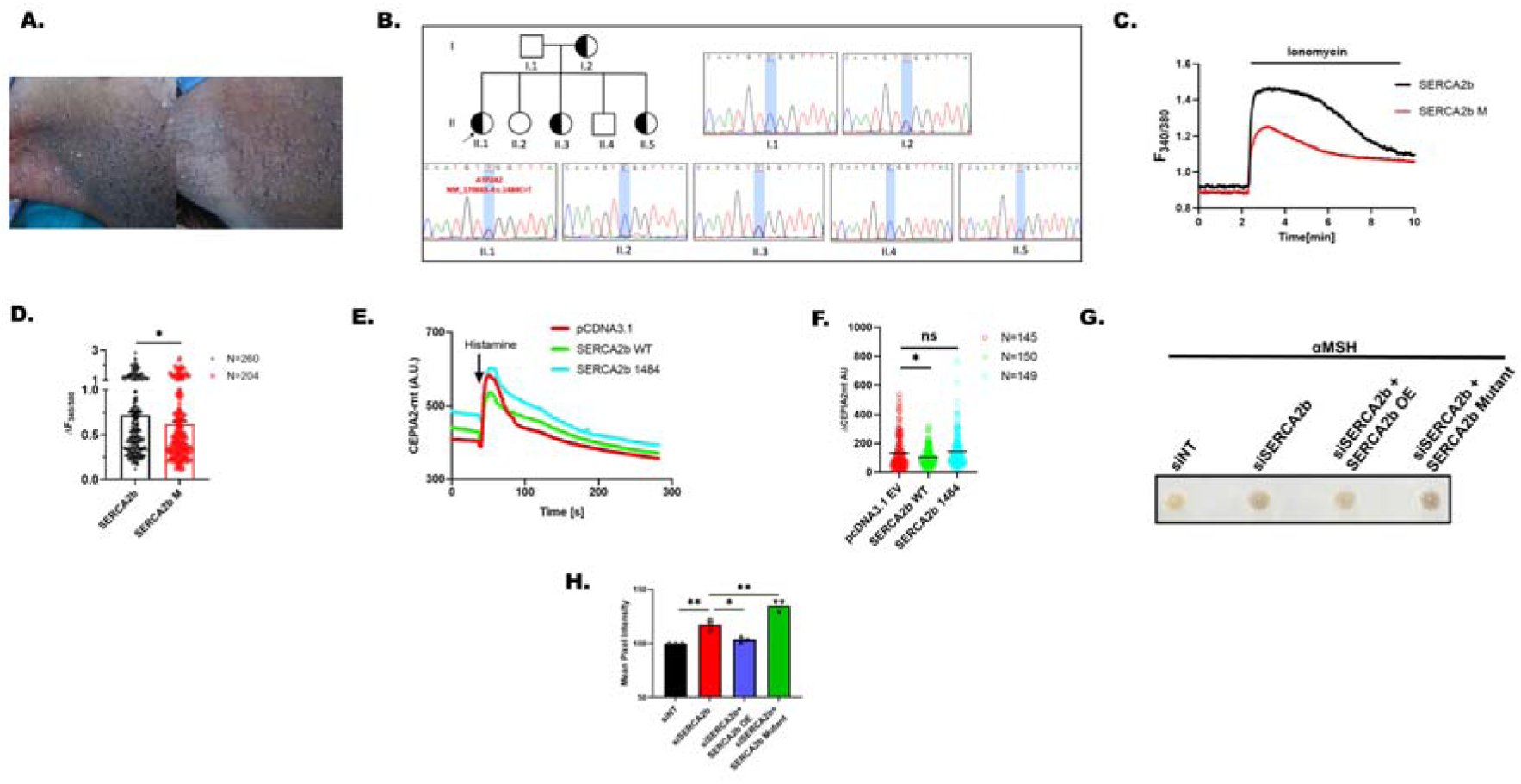
Darier disease associated SERCA2b mutation recapitulates phenotype and leads to reduction in ER Ca^2+^. **(A)** Clinical image showing multiple discrete, warts, hyperkeratotic papules affecting the flexural regions, accompanied by pustules, crusted erosions, and areas of hyperpigmentation. **(B)** Pedigree of the family (Symbols: half-black = affected (heterozygous variant); open = unaffected (wild-type); arrow = index patient) and Sequence chromatogram depicting the identified heterozygous variant (I.2, II.1, II.3 and II.5) and homozygous wild-type allele (I.1, II.2 and II.4) in the family. **(C)** Representative Ca^2+^ imaging trace of WT SERCA2b control and SERCA2b 1484 mutant. Cells were stimulated with 10μM Inomycin in Ca^2+^-free buffer after 48hrs post transfection. **(D**) The extent of ER Ca^2+^ release was calculated from SERCA2b WT Control SERCA2b 1484 mutant, which were imaged from three independent experiments (where “N” denotes total number of cells imaged). **(E**) Representative mitochondrial Ca^2+^ imaging traces of CEPIA2mt in pcDNA control plasmid, SERCA2b WT and SERCA2b 1484 mutant in B16 cells stimulated with 100μM histamine. **(F**) Quantitation of mitochondrial Ca^2+^ uptake by calculating increase in CEPIA2mt signal (ΔCEPIA2mt) in pcDNA control plasmid, WT SERCA2b and SERCA2b 1484 mutant B16 cells stimulated with 100μM histamine where “n” denotes the number of ROIs. **(G**) Representative pellet images of (siNT) control, siSERCA2b, siSERCA2b+SERCA2b OE and siSERCA2b+SERCA2b mutant in B16 cells with αMSH treatment (N=3). **(H**) Quantitation of mean pixel intensity of (siNT) control, siSERCA2b, siSERCA2b+SERCA2b OE and siSERCA2b+SERCA2b mutant in B16 cells with αMSH treatment (N=3). Data presented are mean ± S.E.M. For statistical analysis, unpaired student’s t-test was performed for panel D, and one-way ANOVA followed by Tukey’s post hoc test for panels F and H using Graphpad Prism software. (Here, ns means nonsignificant * p <0.05; ** p < 0.01; ∗∗∗ p < 0.001 and ∗∗∗∗p < 0.0001)

Next, we generated SERCA2b C1484T (S495L) point mutant to examine effect of this mutant on ER Ca^2+^ levels. We measured ER Ca^2+^ levels using FURA2-AM upon overexpression of (WT) SERCA2b, and SERCA2b C1484T mutant in B16 cells. We utilized ionomycin to mobilize ER Ca^2+^ stores into cytosol. We observed significantly lower ER Ca^2+^ stores upon overexpression of SERCA2b C1484T mutant as compared to (WT) SERCA2b (**Fig. 5C and D**).

Further, we examined the effect of SERCA2b C1484T mutant on mitochondrial Ca^2+^ uptake. We performed live cell mitochondrial Ca^2+^ imaging assays on the cells overexpressing empty vector control, WT SERCA2b and SERCA2b C1484T mutant. As shown in **Figure 5E-F**, we observed a decrease in mitochondrial Ca^2+^ uptake upon WT SERCA2b overexpression that was rescued with SERCA2b C1484T mutant overexpression. This data shows that SERCA2b C1484T mutant increases mitochondrial Ca^2+^ uptake.

Next, we asked what happens to αMSH stimulated physiological pigmentation upon silencing of SERCA2b and what is the effect of overexpression of WT SERCA2b and SERCA2b C1484T mutant respectively. We observed increase in pigmentation with siSERCA2b that was decreased upon overexpressing WT SERCA2b, while SERCA2b C1484T mutant lead to increase in pigmentation as compared to siSERCA2b alone (**Fig. 5G and H**).

### 4PBA reverse molecular and phenotypic changes associated with SERCA2b mutation

Next, we employed the SERCA2b C1484T (S495L) point mutant to investigate whether this mutation contributes to the increased pigmentation phenotype observed in patients with Darier disease and to determine if 4PBA can mitigate this pigmentation phenotype. We overexpressed either empty vector control or SERCA2b C1484T mutant or SERCA2b C1484T mutant plus 4PBA treatment in B16 cells and followed (αMSH)-stimulated physiological pigmentation. Interestingly, we observed that overexpression of SERCA2b C1484T mutant results in increase in αMSH stimulated pigmentation as compared to empty vector control and 4PBA treatment results in decrease in SERCA2b C1484T mutant induced hyperpigmentation (**Fig. 6A**). We further quantitated these phenotypic changes by Image J based analysis and observed decrease in SERCA2b C1484T mutant induced hyperpigmentation upon 4PBA treatment (**Fig. 6B**). Further, we examined MERCs and mitochondrial Ca^2+^ uptake downstream of SERCA2b C1484T mutant and SERCA2b C1484T mutant+ 4PBA treatment.

**Figure 6.**
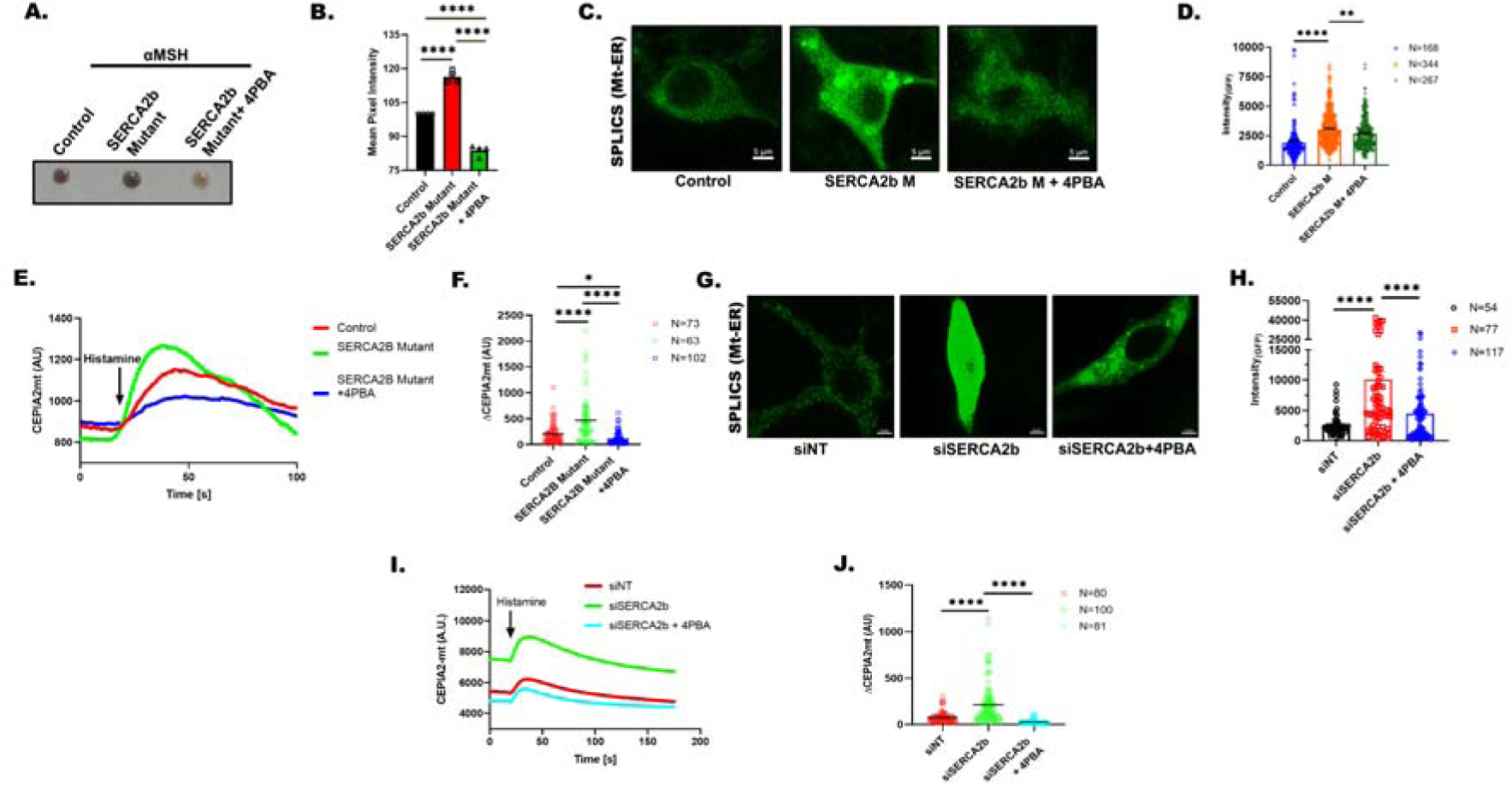
4PBA reverse molecular and phenotypic changes associated with SERCA2b loss of function. **(A)** Representative pellet images of control plasmid, SERCA2b mutant overexpression and SERCA2b mutant overexpression+4PBA in B16 cells with αMSH treatment (N=3). **(B)** Quantitation of mean pixel intensity of control plasmid, SERCA2b mutant overexpression and SERCA2b mutant overexpression+4PBA in B16 cells with αMSH treatment (N=3). **(C)** Representative images of confocal imaging in B16 cells transfected with SERCA2b M and SERCA2b M treated with 1mM 4PBA along with SPLICS Mt-ER Short P2A construct, scale bar, 5µm. **(D)** Bar graph showing the quantification of the GFP intensity in B16 cells transfected with SERCA2b M and SERCA2b M treated with 1mM 4PBA along with SPLICS Mt-ER Short P2A construct, where ‘N’ denotes the number of cells imaged. **(E)** Representative mitochondrial Ca^2+^ imaging traces of control plasmid, SERCA2b mutant overexpression and SERCA2b mutant overexpression+4PBA B16 cells stimulated with 100μM histamine. **(F)** Quantitation of mitochondrial Ca^2+^ uptake by calculating increase in CEPIA2mt signal (ΔCEPIA2mt) in control plasmid, SERCA2b mutant overexpression and SERCA2b mutant overexpression+4PBA B16 cells stimulated with 100μM histamine. **(G)** Representative images of confocal imaging in B16 cells transfected with siSERCA2b and siSERCA2b treated with 1mM 4PBA along with SPLICS Mt-ER Short P2A construct, scale bar, 5µm. **(H)** Bar graph showing the quantification of the GFP intensity in B16 cells transfected with siSERCA2b and siSERCA2b treated with 1mM 4PBA along with SPLICS Mt-ER Short P2A construct, where ‘N’ denotes the number of cells imaged. **(I)** Representative mitochondrial Ca^2+^ imaging traces of CEPIA2mt in siNon-Targeting (siNT) control, siSERCA2b and siSERCA2b+4PBA B16 cells stimulated with 100μM histamine. **(J)** Quantitation of mitochondrial Ca^2+^ uptake by calculating increase in CEPIA2mt signal (ΔCEPIA2mt) in v(siNT) control, siSERCA2b and siSERCA2b+4PBA B16 cells stimulated with 100μM histamine. Data presented are mean ± S.E.M. For statistical analysis, one-way ANOVA followed by Tukey’s post hoc test was performed for panels B, D, F, H and J using Graphpad Prism software. (Here, * p <0.05; ** p < 0.01; ∗∗∗ p < 0.001 and ∗∗∗∗p < 0.0001)

We utilized split GFP-based contact site sensor SPLICS_S_ (Mt-ER) to measure ER-mitochondrial distance upon overexpression of SERCA2b C1484T mutant and SERCA2b C1484T mutant along with 4-PBA treatment (Cieri et al., 2017). We observed increase of short-distance MERCs upon overexpression of SERCA2b C1484T mutant as compared to control which was rescued with 4PBA in SERCA2b C1484T mutant+ 4PBA overexpressing cells as compared to SERCA2b C1484T mutant (**Fig. 6C and D**). This data suggests that ER-mitochondrial distance decreases or ER-mitochondria come closer upon overexpression of SERCA2b C1484T mutant. Next, we asked whether these increased ER-mitochondrial contacts could increase mitochondrial Ca^2+^ downstream of SERCA2b C1484T mutant overexpression.

Next, we examined the effect of SERCA2b C1484T mutant and SERCA2b C1484T mutant+ 4PBA on mitochondrial Ca^2+^ uptake. We performed live cell mitochondrial Ca^2+^ imaging assays on the cells transfected either with empty vector control or SERCA2b C1484T mutant alone or SERCA2b C1484T mutant along with 4-PBA treatment. As presented in **Figure 6E-F**, the increase in mitochondrial Ca^2+^ uptake in SERCA2b C1484T mutant condition is rescued upon 4-PBA treatment **(Fig 6E-F)**. This data shows that SERCA2b C1484T mutant increases mitochondrial Ca^2+^ uptake and rescued upon 4-PBA treatment.

Further, we asked whether 4-PBA treatment can rescue ER-mitochondrial contacts and mitochondrial Ca^2+^ uptake downstream of SERCA2b silencing. We utilized split GFP-based contact site sensor SPLICS_S_ (Mt-ER) to measure ER-mitochondrial distance upon transfection with siNT control, siSERCA2b alone and siSERCA2b along with 4-PBA treatment (Cieri et al., 2017). We observed decrease of MERCs upon siSERCA2b + 4-PBA treatment as compared to SERCA2b silenced cells (**Fig. 6G and H**). This data suggests that ER-mitochondrial distance decreases or ER-mitochondria come closer upon SERCA2b silencing and this phenomenon is rescued by 4-PBA treatment. Next, we asked whether these increased ER-mitochondrial contacts could increase mitochondrial Ca^2+^ downstream of SERCA2b silencing. Further, we examined the effect of 4-PBA mediated alleviation of ER stress on mitochondrial Ca^2+^ uptake. We performed live cell mitochondrial Ca^2+^ imaging assays on the cells transfected either with siSERCA2b alone or siSERCA2b along with 4-PBA treatment. The increase in mitochondrial Ca^2+^ uptake in siSERCA2b condition is rescued upon 4-PBA treatment **(Fig 6 I-J)**. This data shows that SERCA2b silencing driven ER stress increases mitochondrial Ca^2+^ uptake and it is rescued upon 4-PBA treatment.

### 4PBA inhibits physiological pigmentation

Next, we investigated the potential of targeting ER stress in controlling pigmentation. We used 4PBA to determine the significance of alleviating ER stress in modulating melanogenesis. We utilized *in vivo* zebrafish model system and treated 2 dpf (days post fertilization) zebrafish embryos with 4PBA. Phenotypically, we observed significant decrease in the pigmented black-colored melanophores in 4PBA treated morphants as compared to the control embryos 48h post treatment (**Fig. 7A**). We quantitated this decrease in pigmentation by Image J based analysis (**Fig. 7B**). Next, we used αMSH for inducing pigmentation in B16 cells and co-treated cells with 4PBA to examine contribution of ER stress in αMSH stimulated pigmentation. Interestingly, we observed that 4PBA treatment results in decrease in αMSH stimulated physiological pigmentation (**Fig. 7C**). We further quantitated these phenotypic changes by Image J based analysis and observed decrease in αMSH induced melanogenesis upon 4PBA treatment (**Fig. 7D**). This data suggests that 4PBA can be used for modulating pigmentation.

**Figure 7.**
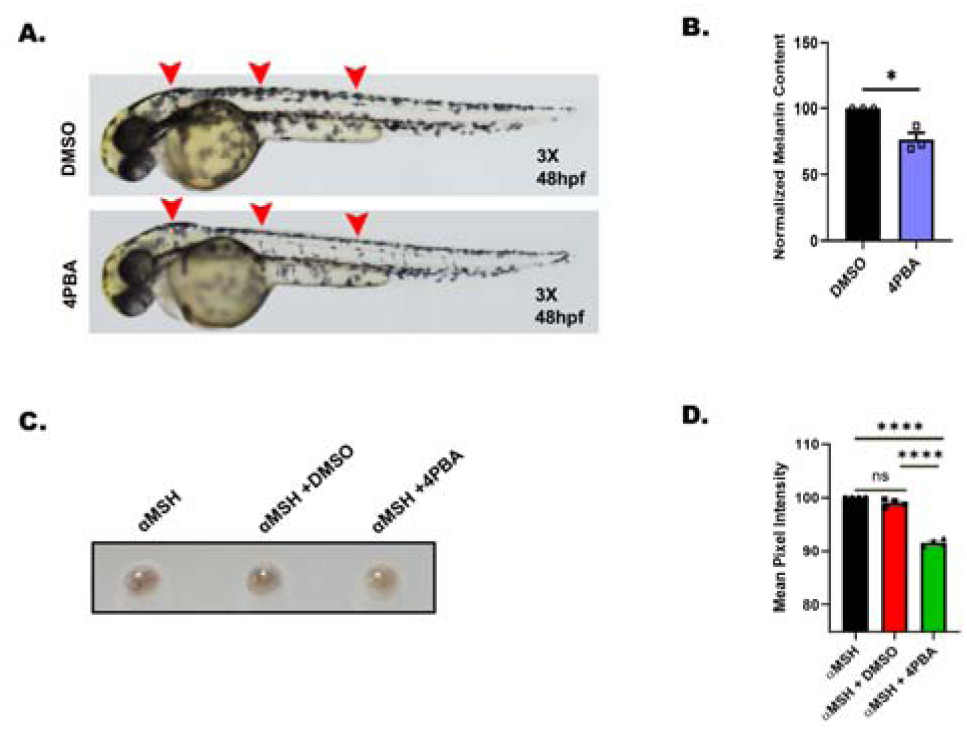
4PBA inhibits physiological pigmentation. **(A)** Representative bright-field images of wild-type zebrafish embryos treated with either DMSO control or 4PBA at 48hpf (hours post fertilization) (N=3, 20 embryos in each “N”). **(B)** Melanin content estimation of zebrafish embryos treated with either DMSO control or 4PBA at 48hpf (hours post fertilization) (N=3, 20 embryos in each “N”). **(C)** Representative pellet images of B16 cells treated with αMSH, αMSH+DMSO and αMSH+4PBA (N=3). **(D)** Quantitation of mean pixel intensity of B16 cells treated with αMSH, αMSH+DMSO and αMSH+4PBA (N=3). Data presented are mean ± S.E.M. For statistical analysis, one sample t test was performed for panels B using Graphpad Prism software. (Here, * p <0.05; ** p < 0.01; ∗∗∗ p < 0.001 and ∗∗∗∗p < 0.0001)

**Figure 8.**
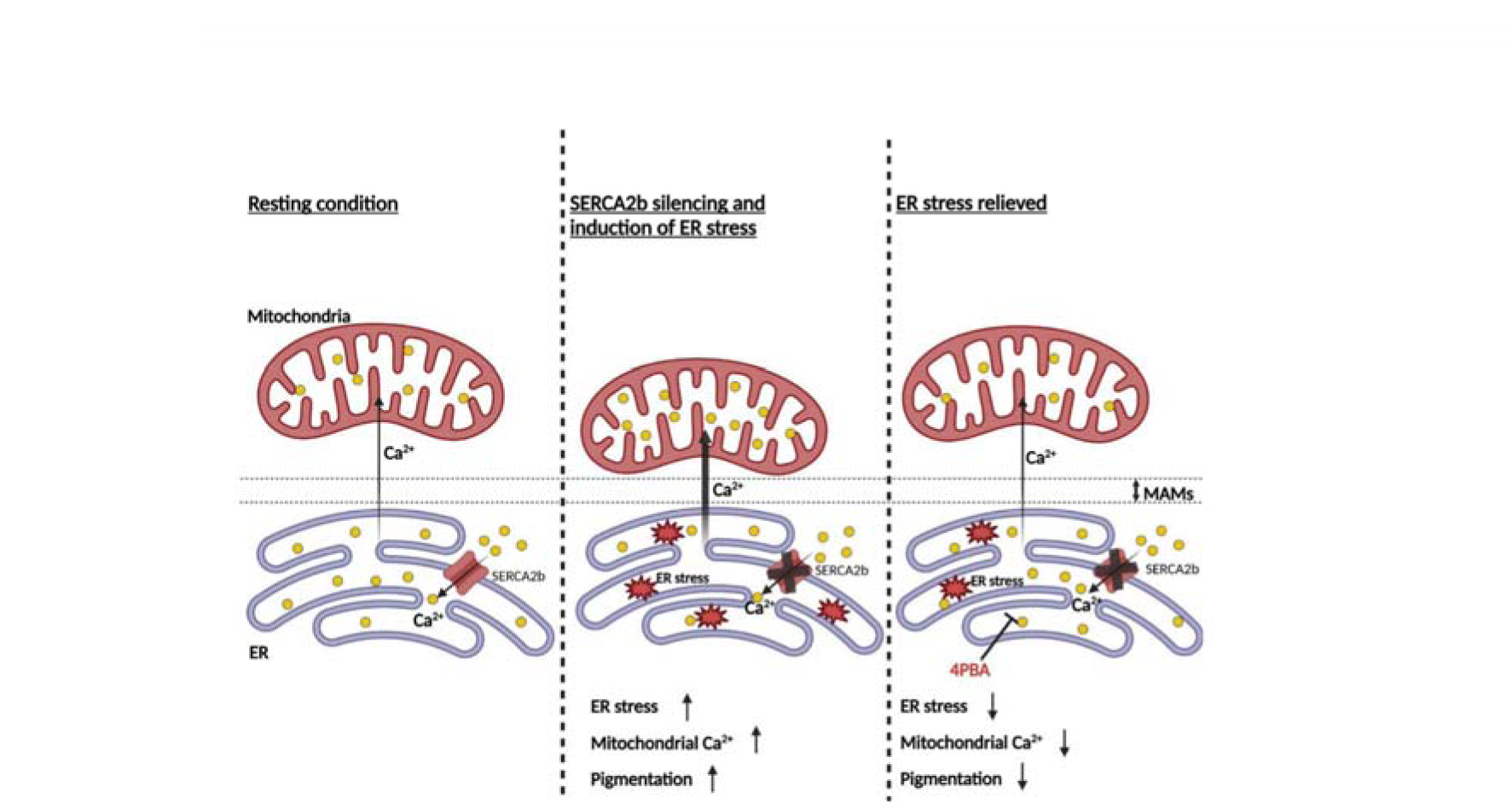
SERCA2b acts as a negative regulator of pigmentation by inducing adaptive ER stress and enhancing mitochondrial calcium uptake.

## Discussion

ER and Mitochondrial Ca^2+^ signaling is emerging as a critical regulator of skin pigmentation. Our earlier work revealed that physiological melanogenic stimuli αMSH stimulates ER Ca^2+^ efflux that leads to activation of ER Ca^2+^ sensor Stromal interaction molecule1 (STIM1) (Motiani et al., 2018). STIM1 in turn induces pigmentation by augmenting cAMP generation (Motiani et al., 2018). Recently, we demonstrated that αMSH enhances STIM1 transcription via master pigmentation transcription factor MITF (Tanwar, Sharma, et al., 2022). Interestingly, in tanned human skin samples STIM1 expression shows a very strong co-relation with the MITF levels thereby highlighting the physiological relevance of this signaling module (Tanwar, Sharma, et al., 2022). Further, literature suggests that UV-rays stimulate ER Ca^2+^ efflux and that in turn contributes to pigmentation (Ahuja et al., 2024; Bellono et al., 2014; Stanisz et al., 2012). Taken together, ER Ca^2+^ signaling plays an important role in physiological pigmentation.

In our study, we identify SERCA2b as a negative regulator of pigmentation. Our data establish SERCA2b as a key determinant of pigmentation by maintaining Ca^2+^ levels in the ER. Here, we show that SERCA2b silencing results in decrease in ER Ca^2+^ levels. This in turn leads to enhanced pigmentation, adaptive ER stress (**Figure 1 and 3**) and enhanced mitochondrial Ca^2+^ uptake (**Figure 4**). SERCA2b silencing triggers activation of adaptive ER stress markers as Bip, sXBP1 and tXBP1 (**Figure 3**). This adaptive ER stress reorganizes ER-mitochondria contact sites (MAMs), thereby increasing mitochondrial Ca²L uptake (**Figure 4**). We recently revealed an evolutionarily conserved role of mitochondrial Ca^2+^ uptake in vertebrate pigmentation (Tanwar et al., 2024). In our study, we showed that enhanced mitochondrial Ca²L uptake via MCU promotes pigmentation via upregulating keratin filament expression, which further potentiates melanogenesis by regulating melanosome biogenesis and maturation (Tanwar et al., 2024). Our data therefore suggest that SERCA2b loss-of-function increases pigmentation via ER stress-mediated remodeling of ER-mitochondrial distance and subsequent increase in mitochondrial Ca²L uptake.

We observe that SERCA2b loss of function augments mitochondrial Ca²L uptake through ER–mitochondrial tethering, suggesting that SERCA2b and MCU may operate along a shared signaling axis to fine-tune mitochondrial Ca²L dynamics and melanogenesis. Thus, while MCU acts as a positive regulator of pigmentation by enhancing mitochondrial Ca²L uptake, SERCA2b functions as a negative regulator that restrains this pathway by maintaining ER Ca²L homeostasis. The opposing activities of SERCA2b and MCU together define a Ca²L-dependent regulatory network that balances ER stress, mitochondrial Ca²L uptake, and pigmentation.

Further, alleviation of ER stress in siSERCA2b condition with 4-PBA rescues pigmentation phenotype as well as mitochondrial Ca^2+^ uptake (**Figure 3 and 6**). Thus, adaptive ER stress emerges as a potential therapeutic target for the treatment of hyperpigmentary disorders. Given the established clinical safety of 4-PBA, repurposing it or similar ER stress modulators could represent a feasible therapeutic approach for Darier’s disease as well as other pathological hyperpigmentary disorders. Moreover, our study suggests that fine-tuning ER-mitochondrial Ca²L signaling could serve as a novel strategy to modulate pigmentation.

Collectively, our study uncover a previously unknown SERCA2b–ER stress–mitochondrial Ca²L axis as a central regulator of pigmentation. SERCA2b acts as a negative regulator of pigmentation by maintaining ER Ca^2+^ levels and suppressing adaptive ER stress. SERCA2b loss-of-function-promotes ER-mitochondrial coupling, thereby increasing mitochondrial Ca²L uptake, and melanogenesis. Future studies should aim to explore how other ER stress pathways (e.g., PERK, IRE1, ATF6) interface with melanogenic transcriptional networks and whether similar mechanisms operate in UV-induced pigmentation or other pigmentary disorders. In summary, our study identify SERCA2b as a negative regulator of pigmentation and provides a mechanistic and therapeutic framework for managing pigmentary disorders associated with ER Ca²L dysregulation.

## Authors Contribution

Akshay Sharma: Methodology, Investigation, Visualization, Formal analysis. Suman Saurav: Methodology, Investigation, Visualization, Formal analysis, Writing-Original draft preparation. Preeti Sharma: Investigation, Visualization, Formal analysis. Anushka Agrawal: Investigation, Visualization, Formal analysis. Nutan Sharma: Investigation. Gyan Ranjan: Methodology, Investigation, Visualization, Formal analysis. Divya Bhalla: Visualization, Formal analysis. Deepika Pandhi: Investigation, Visualization, Formal analysis. Vamsi K Yenamandra: Investigation, Visualization, Formal analysis. Jyoti Tanwar: Methodology, Investigation, Visualization, Formal analysis, Writing-Original draft preparation. Rajender K Motiani: Conceptualization, Supervision, Writing-Original draft preparation, Reviewing and Editing, Project administration, Funding acquisition.

## Acknowledgements

This work was supported by the DBT/Wellcome Trust India Alliance Fellowship (IA/I/19/2/504651). RKM also acknowledges funding support from RCB Institutional Core funding (R25226). The authors thank members of the Motiani laboratory for the critical discussions. SS and NS acknowledge their Junior and Senior Research Fellowship from CSIR India and DBT India respectively.

